# Targeted activation of microglial PPARδ reprograms immunometabolism and enhances insulin sensitivity in diet-induced obesity

**DOI:** 10.1101/2024.11.18.624062

**Authors:** Han Jiao, Fernando Cázarez-Márquez, Shanshan Guo, Andries Kalsbeek, Gertjan Krammer, Chun-Xia Yi

## Abstract

Microglia play a crucial role in maintaining neuronal health through phagocytosis, a function that becomes compromised during diet-induced obesity and is associated with altered lipid metabolism. Previous research demonstrated that disrupting lipid metabolism in microglia, such as through lipoprotein lipase deficiency, impairs their phagocytic function and exacerbates obesity, glucose dysregulation, and hypothalamic neuron dysfunction. This study investigated whether enhancing lipid metabolism via peroxisome proliferator-activated receptor delta (PPARδ) activation could counteract obesity-related metabolic disturbances. Thermal proteome profiling identified GW0742 as the most potent PPARδ ligand among those tested. GW0742 enhanced microglial phagocytosis, reduced inflammation, and shifted energy metabolism towards glycolysis over oxidative phosphorylation. Targeted delivery of GW0742 using nanoparticles (NPs-GW0742) to microglia in the mediobasal hypothalamus of obese rats significantly improved insulin sensitivity without affecting body weight or food intake. Enhanced microglial activation was evidenced by increased soma size and coverage. These findings underscore the importance of microglial lipid metabolism in systemic glucose regulation and highlight the potential of PPARδ-targeted therapies to mitigate hypothalamic inflammation and improve metabolic health in obesity.

## Introduction

To sustain a healthy neuronal microenvironment, microglia, the predominant innate immune cells of the brain, play a critical role through their phagocytic function, which involves the clearance of cell debris and pathogens [24, 30]. Our previous work has demonstrated that in diet-induced obesity, particularly following a high-fat diet (HFD), the increased metabolic stress leads to an accumulation of cellular debris and metabolic waste products from neurons and surrounding cells. This condition necessitates heightened microglial phagocytic and immune activity, particularly in the hypothalamus [13–16, 33]. These processes are tightly linked to increased lipid utilization and enhanced lipoprotein lipase (LPL) activity in microglia [16]. Importantly, microglia deficient in LPL exhibit reduced lipid uptake, impaired immune responses, diminished phagocytic capacity in microglia, and an accelerated rate of weight gain when subjected to an HFD, in association with worsened glucose metabolism and impairments of pro-opiomelanocortin (POMC) neurons that control food intake, energy expenditure and glucose metabolism [9]. Therefore, in the context of diet-induced obesity, effective lipid utilization by microglia is essential for maintaining their immune and phagocytic functions. Enhancing microglial lipid metabolism could thus potentially yield anti-obesity effects.

One promising strategy for augmenting lipid metabolism in cells is the activation of peroxisome proliferator-activated receptor (PPAR) pathways, known for their central role in promoting fatty acid oxidation [32, 36]. The PPAR family comprises three isoforms-PPARα, PPARγ, and PPARδ [4]-all of which contribute to the regulation of lipid and glucose metabolism, thereby supporting energy homeostasis [38]. In the central nervous system, the distribution and expression of PPARs are region-and cell-type specific. PPARα is predominantly found in the hippocampus, cerebellum, olfactory bulb, and retina, whereas PPARγ displays limited distribution [6, 28], PPARδ, however, shows high expression in the mediobasal hypothalamus (MBH), implicating it in energy homeostasis [28, 39, 40]. On a cellular level, PPARs are present in neurons, while astrocytes mainly express PPARα and PPARγ [37], Microglia notably express the highest levels of PPARδ [43, 44]. Prior studies have shown that PPARδ-deficient mice on an HFD exhibit impaired energy uncoupling and are prone to obesity [36]. Additionally, microglial PPARδ deficiency has been linked to reduced proliferation and heightened proinflammatory cytokine production following LPS exposure [10]. However, the specific impact of targeted activation of microglial PPARδ on immune and metabolic function, and its broader implications for neural circuits that regulate feeding behavior, body weight, and glucose homeostasis, remains poorly understood.

In this study, we aim to elucidate the pharmacodynamics of distinct PPARδ ligands in microglia, assessing both their on-target and off-target effects [36], through thermal proteome profiling (TPP)-a method leveraging the principle that ligand-bound proteins exhibit increased resistance to heat-induced denaturation [9]. We will evaluate two novel PPARδ ligands, GNF-0242 and GNF-8501 [9], alongside the established PPARδ agonist GW0742 [34]. The *in vitro* effects of these ligands on microglial immunometabolic function will be characterized, and the most potent compound will be delivered specifically to microglia in the MBH of HFD-induced obese rats to investigate its effects on food intake, body weight, and glucose metabolism. To specifically deliver a PPARδ ligand into microglia in the MBH, we have established a nanoparticles (NPs)-mediated delivery method. This method is developed based on our previous study on using NPs to deliver siRNA into microglia in the hypothalamus for genetic modification [18].

## Results

### Thermal proteome profiling of GNF-0242, GNF-8501, GW0742

Thermal proteome profiling (TPP) is a robust method for assessing the thermal stability and abundance of proteins, enabling the unbiased identification of direct and indirect drug targets [11, 31]. This technique offers significant advantages over traditional proteomic approaches, including high throughput, sensitivity, precision, and the ability to provide comprehensive insights into complex biological systems [3]. Ligand binding typically enhances the thermal stability of target proteins compared to their unbound state [25]. We utilized TPP to evaluate proteomic changes following the incubation of microglia with the PPARδ ligands GNF-0242, GNF-8501, and GW0742 (**Fig. 1**). Proteins exhibiting altered thermal stability compared to the vehicle control were identified as potential off-targets of the ligands, indicating reduced specificity (**Fig. 1A**). Our analysis revealed that, compared to control treatment, more than 2,000 proteins displayed significant changes in thermal stability across the three PPARδ agonists (**Fig. 1B**). Specifically, we detected significant melting point shifts in 75, 82, and 27 proteins for GNF-0242, GNF-8501, and GW0742, respectively (**Fig. 1C**). Overall, these findings suggest that GNF-0242 and GNF-8501 interact with a broader range of proteins than GW0742, indicating a higher potential for off-target effects, consistent with the observations from immune challenge studies (**Fig. 1D**).

**Figure 1.**
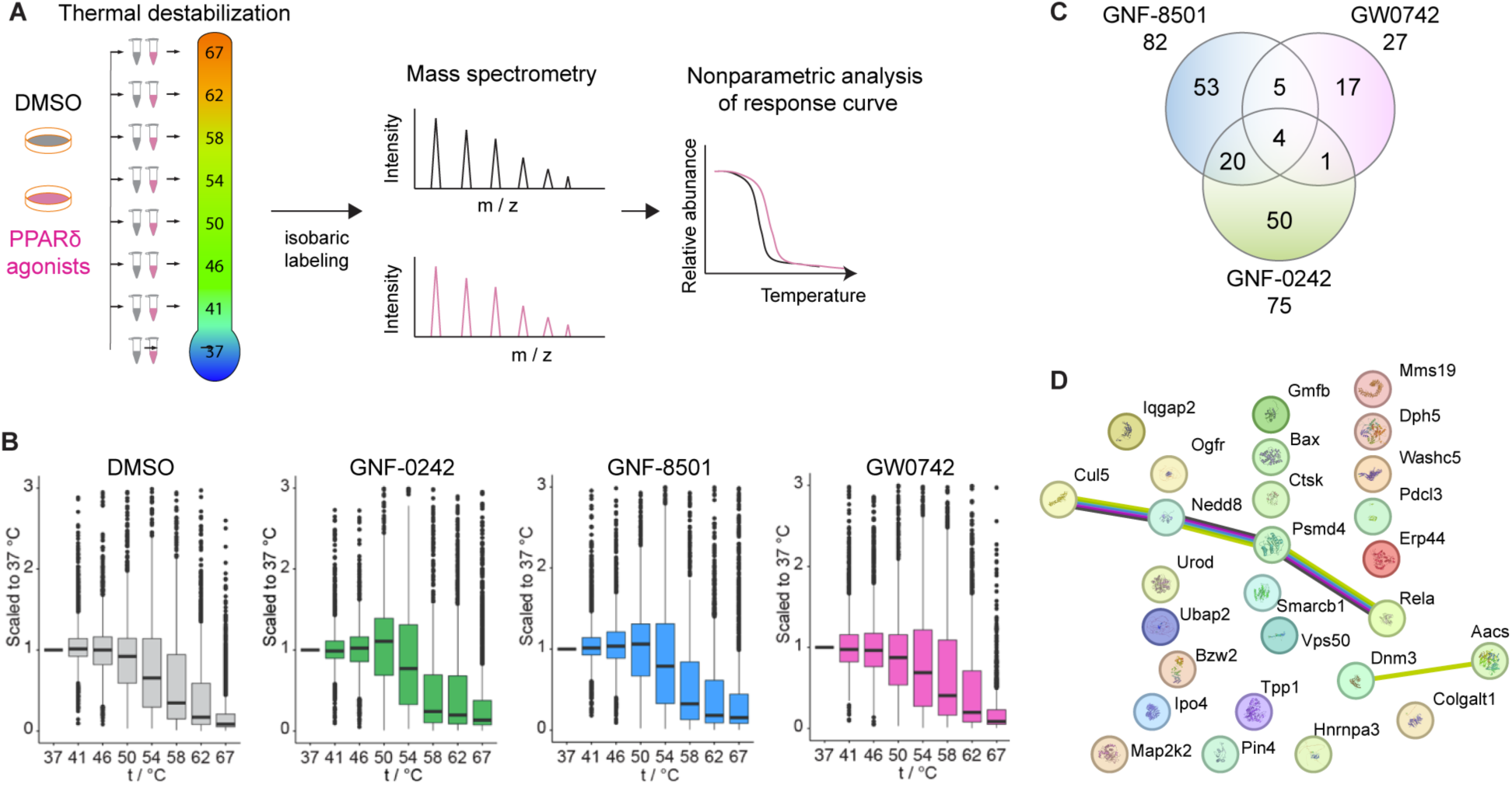
Thermal Proteome Profiling of PPARδ agonists in microglial cells. A, Overview of the experimental setup. PPARδ agonists and vehicle (DMSO) treatment and 8-temperature destabilization were performed. B, Distribution of TPP data after scaling each protein to the abundance signal in the lowest temperature condition (37 °C). The solid line in the box plots represents the median, box limits show the interquartile range (IQR) and its whiskers 1.5 x IQR. Outliers are represented as block dots. C, Venn diagram representing the number of proteins with shifted temperatures, compared to control treatment for each of the three tested agonists-GNF-0242, GNF-8501 or GW0742. D, Protein networks with increased temperature induced by GW0742.

Among the proteins with altered melting shared by all three agonists, four were identified: RELA, CTSK, and MAP2K2 showed increased temperatures, while AACS displayed decreased temperatures (**Fig. 1C**). For GW0742 specifically, 27 proteins exhibited increased temperatures. Functional enrichment analysis indicated that four of these proteins-CLU5, NEDD8, PSMD4, and RELA-are part of a network associated primarily with cellular stress response and protein degradation (**Fig. 1D**). These data underscore the potential pathways modulated by GW0742, highlighting its more selective profile compared to GNF-0242 and GNF-8501.

### GW0742 demonstrates superior potency among the three PPARδ agonists

Given the involvement of the pathways identified in Fig. 1C and the pivotal neuroprotective role of microglial phagocytosis, we assessed the effects of the three PPARδ agonists on microglial phagocytic activity. Our analysis revealed that only GW-0742 significantly enhanced microglial phagocytic function compared to the DMSO control (Fig. 1A and B). HFD exposure is known to compromise the integrity of the intestinal epithelial barrier, leading to increased intestinal permeability [22]. This disruption can elevate circulating lipopolysaccharide (LPS) levels, contributing to peripheral tissue inflammation [21]. To further investigate the influence of PPARδ agonists on microglial immune responses, BV-2 cells treated with PPARδ agonists and control cells were challenged with 100 ng/mL LPS (**Fig. 2, C–F**). LPS treatment markedly increased the expression of pro-inflammatory cytokines *Tnfα* and *Il6* as well as the oxidative stress marker *iNos/Nos2* (**Fig. 2, C, D, F**). Among the tested agonists, only GW-0742 effectively attenuated these pro-inflammatory responses (**Fig. 2, C, D, F**). Moreover, LPS exposure reduced the expression of the anti-inflammatory cytokine *Il10*. Notably, GW0742 treatment showed a trend toward upregulating *Il10* expression, suggesting its potential role in promoting an anti-inflammatory response (**Fig. 2E**). These findings collectively indicate that GW0742 is more effective than GNF-0242 and GNF-8501 in enhancing microglial phagocytosis and modulating the immune response, highlighting its superior potency as a PPARδ agonist.

**Figure 2.**
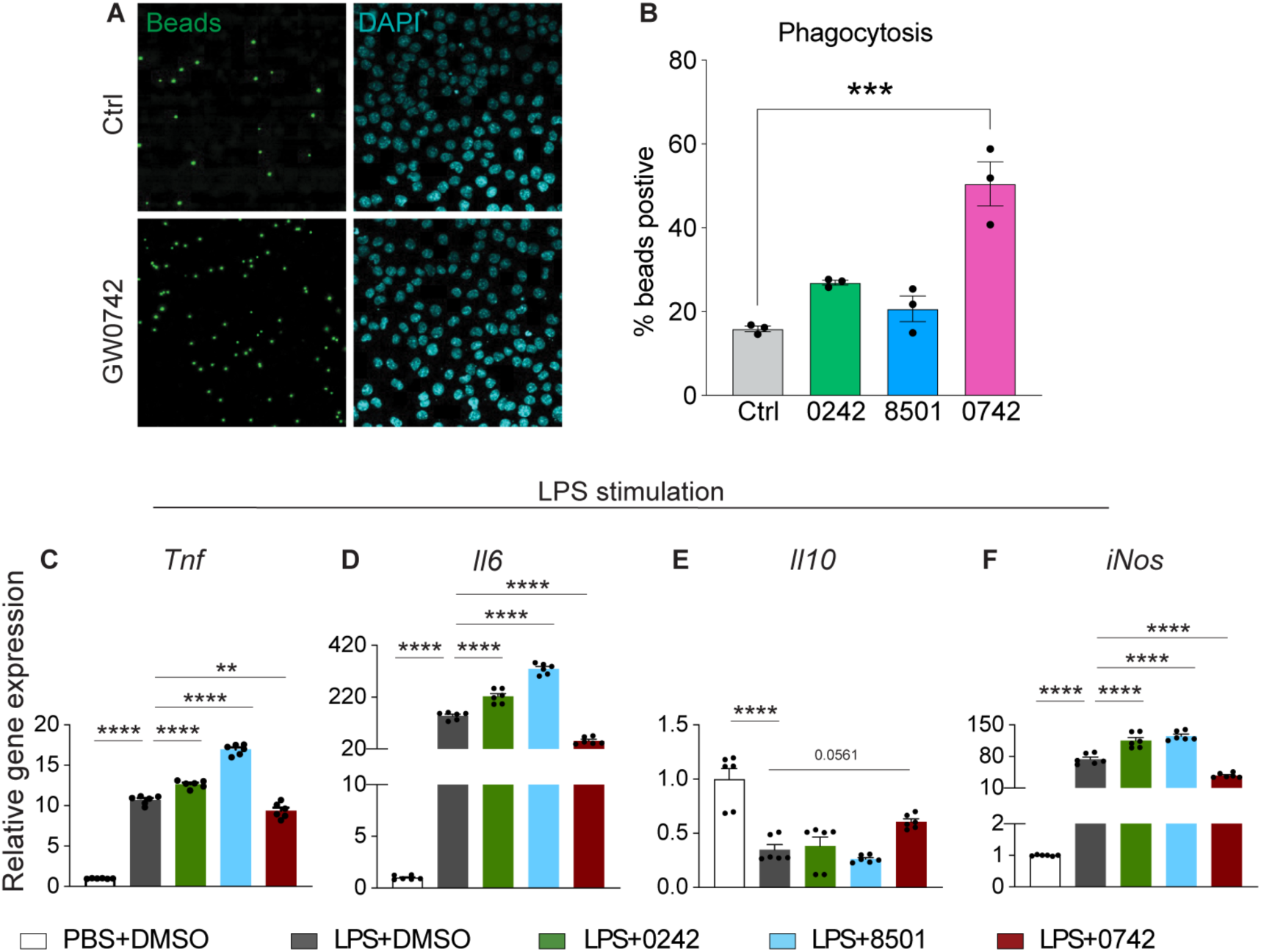
The effect of PPARδ agonists on LPS induced inflammatory responses in microglia cells. A, The uptake of fluorescent beads by primary microglia treated with DMSO or PPARδ agonists. B, The uptake quantification of microspheres. C, Relative gene expression of cytokines (Tnf, Il6, Il10) and oxidative stress-related gene iNos in BV2 microglia treated with PPARδ agonists or DMSO together with 100 ng/mL LPS or PBS treatment for 24 h. Data are presented as means ± SEM and statistical significance was determined using Unpaired t test.

### GW-0742 modulates microglial energetic metabolism

Microglial immune function is closely linked to their metabolic state, an interaction known as immunometabolism, which is essential for regulating inflammatory responses, maintaining neuronal homeostasis, and supporting overall brain function [5]. To investigate the effects of GW0742 on microglial metabolic activity, microglia were treated with 1 μM GW0742 or DMSO (vehicle control) for 24 hours, followed by assessments of oxygen consumption rate (OCR) and extracellular acidification rate (ECAR) using a Seahorse XFe96 analyzer to measure mitochondrial respiration and glycolysis, respectively. The traces for OCR and ECAR and the analyzed metabolic parameters are shown in **Fig. 3A** and **Fig. 4A**, with corresponding raw data in **Fig. 3B** and **Fig. 4B**. GW0742 treatment significantly reduced mitochondrial respiration (**Fig. 3C-E**), primarily due to a decrease in ATP-linked respiration, while the proton leak remained unchanged (**Fig. 3F-H**). Furthermore, GW0742 reduced maximal substrate oxidation capacity (**Fig. 3I, 3J**), while coupling efficiency showed no change (**Fig. 3K**). These findings indicate that GW0742 decreases ATP production by inhibiting oxidative phosphorylation, suggesting that microglia require less energy for immune responses under GW0742 treatment.

**Figure 3.**
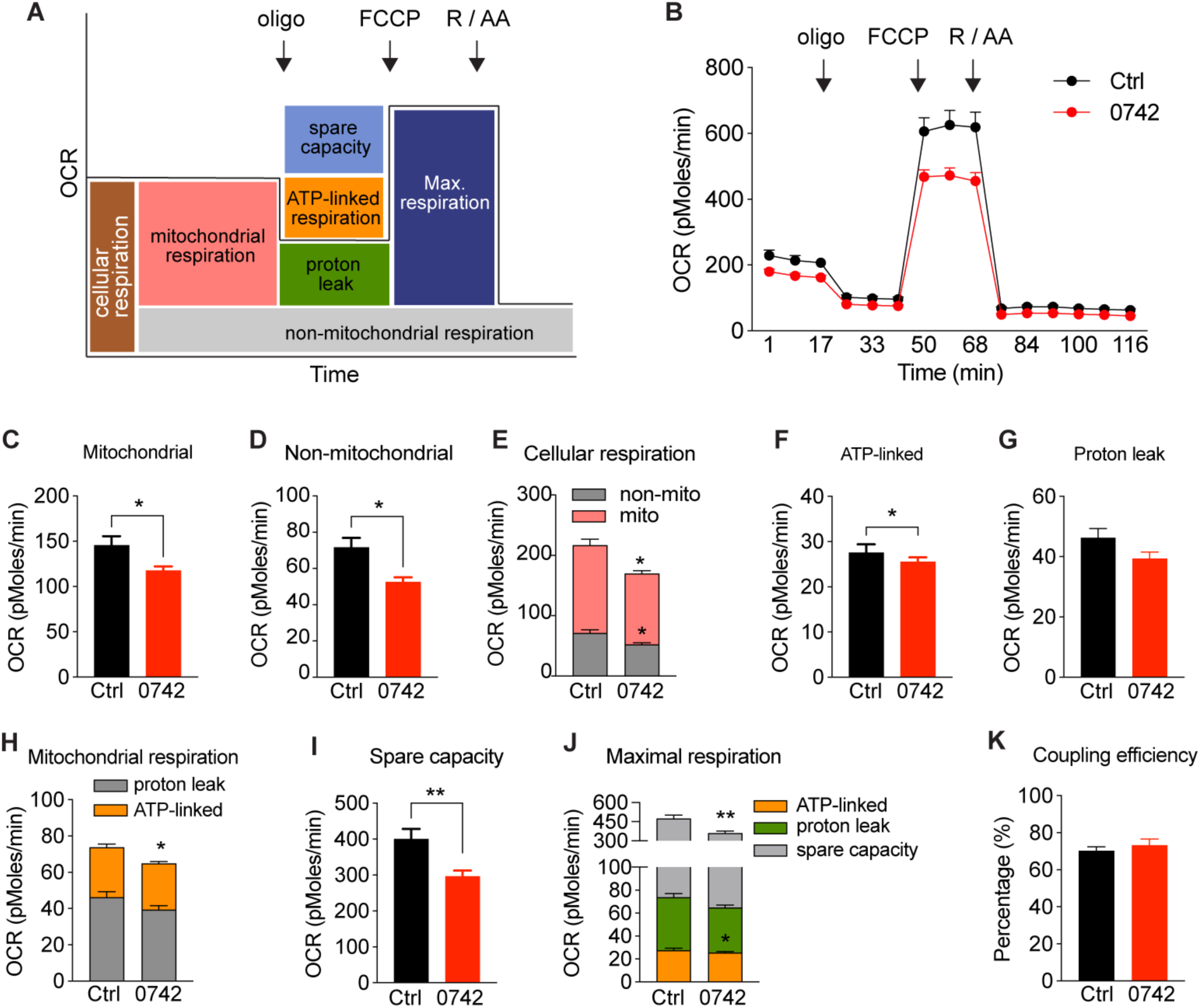
GW0742 reduces microglial ATP-linked respiration without changing coupling efficiency. A, Scheme defining the oxygen-consuming rate (OCR) processes of cells. B, OCR over time in microglial cells treated with GW0742 or vehicle (Ctrl, DMSO). C-E, Cellular respiration was dissected into mitochondrial and nonmitochondrial respiration using the electron transfer chain inhibitors, rotenone and antimycin A. F-H, Mitochondrial respiration was further dissected into ATP-linked respiration and proton leak using the ATP-synthase inhibitor, oligomycin. I-J, Maximal respiration of cells after addition of the chemical uncoupler, FCCP, and after subtracting nonmitochondrial respiration rates. The portion of spare respiratory capacity was determined by subtracting basal respiration from maximal respiration rates. K, Mitochondrial coupling efficiencies were calculated as the fraction of mitochondrial oxygen consumption that is sensitive to oligomycin, reflecting the fraction used to drive ATP synthesis. Data are presented as means ± SEM and statistical significance was determined using Unpaired t test.

**Figure 4.**
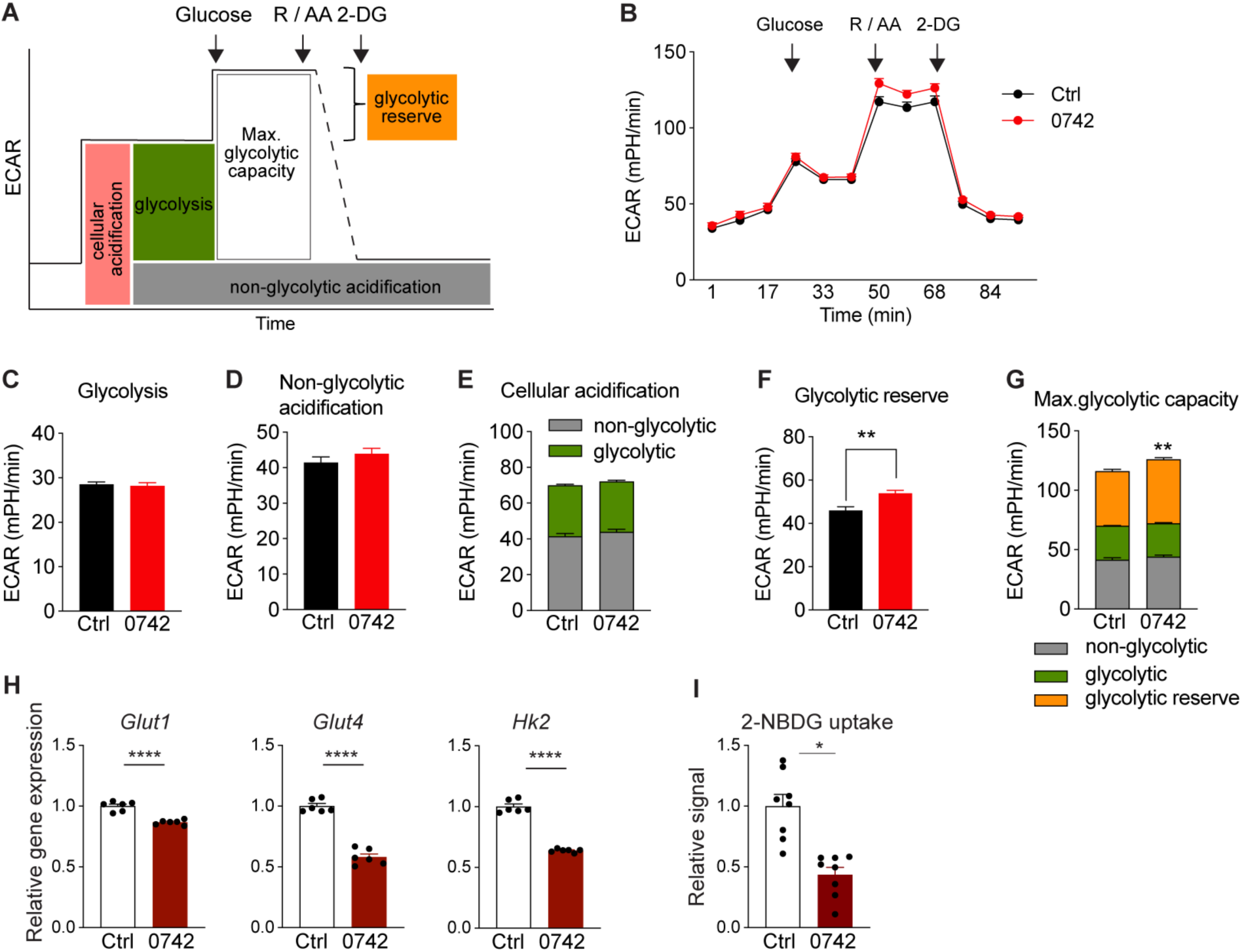
GW0742 reduces microglial glycolytic reserve capacity. A, Scheme defining the cellular extracellular acidification rate (ECAR) process. B, The ECAR over time in BV2 microglial cells treated with GW0742 or vehicle (Ctrl, DMSO). C-E, Cellular acidification was dissected into glycolysis and non-glycolytic acidification. F, G, Glycolytic reserve and maximal glycolytic capacity. H, The expression of glucose metabolism gene Glut1, Glut4 and Hk2. I, 2-NBDG glucose uptake of GW0742 treated group compared to vehicle control group. Data are presented as means ± SEM and statistical significance was determined using Unpaired t-test.

In contrast, while GW0742 treatment did not change basal glycolysis (**Fig. 4A-E**), it increased the maximal glycolytic capacity (**Fig. 4F, 4G**), attributed to an enhanced glycolytic reserve (**Fig. 4F**). This suggests that GW0742 bolsters microglia’s capacity to respond to increased energy demands. Additionally, GW0742 reduced the expression of key enzymes involved in glucose metabolism, such as *Hk2*, *Glut1*, and *Glut4* (**Fig. 4H**), supporting a shift away from glucose utilization in the citric acid cycle. To confirm these observations, we measured glucose uptake using 2-NBDG and found a significant reduction in glucose uptake in GW0742-treated microglia compared to controls (**Fig. 4I**). Collectively, these results indicate that GW0742 shifts microglial energy metabolism, reducing reliance on oxidative phosphorylation and enhancing the capacity for glycolytic response, potentially optimizing energy use to support immune functions.

### Synthesis and characterization of NPs-GW0742 for microglial targeting *in vitro* and *in vivo*

To achieve targeted delivery of GW0742 to microglia within the MBH, we developed a nanoparticle (NP)-based delivery system. These GW0742-encapsulated nanoparticles (NPs-GW0742) were synthesized using amphiphilic mPEG-PLGA monomers (**Fig. 5A**). The resulting NPs-GW0742 had an average diameter of 197.8 nm with a polydispersity index (PDI) of 0.185, indicating uniform size distribution (**Fig. 5B**). The surface charge of the NPs-GW0742 was measured at-12.1 mV (**Fig. 5C**). The morphology and structural integrity of these NPs were confirmed via transmission electron microscopy (TEM), which revealed spherical and uniform particles (**Fig. 5D**). To assess the cellular uptake of these NPs, we conducted a time-course study in microglial cultures using NPs encapsulated with the fluorescent dye rhodamine B (NPs-RhoB). Over a 24-hour period, the RhoB fluorescence intensity per cell increased steadily, demonstrating efficient and sustained NP uptake by microglia (**Fig. 5E**). To verify targeted *in vivo* uptake, we injected NPs-RhoB into the MBH of rat brains and observed the distribution via immunofluorescence staining after 4 hours. The results showed that NPs-RhoB selectively accumulated in microglia, while no significant RhoB signal was detected in astrocytes or neurons (**Fig. 5F-H**).

**Figure 5.**
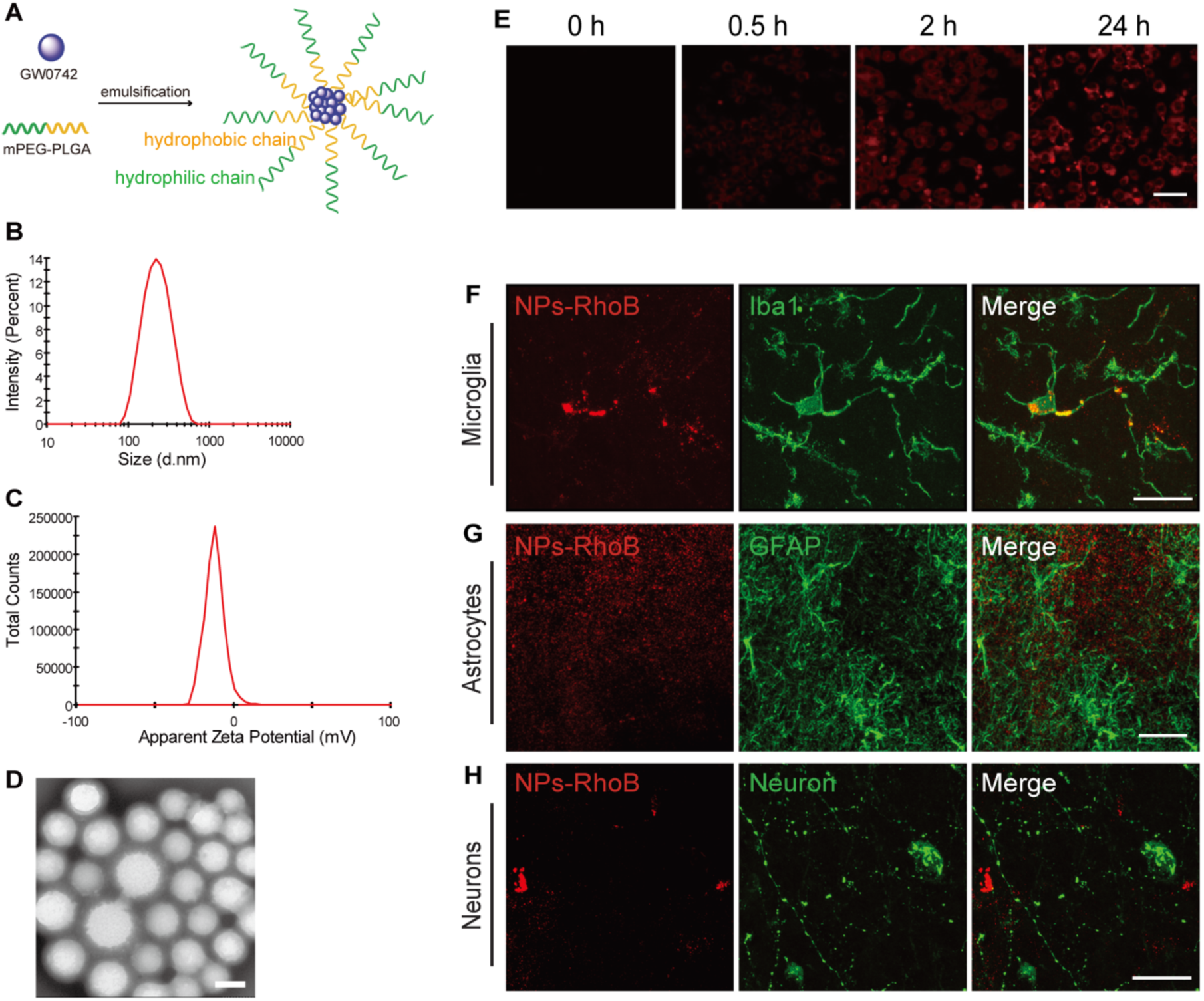
Synthesis and characterization of NPs-GW0742 uptake *in vitro* and *in vivo*. A, Schematic diagram of the synthesis of NPs-GW0742 nanomaterials. B, Particle size distribution and C, surface charge and D, transmission electron microscope (TEM) image of NPs-GW0742. Scale bar in D, 200 nm. E, NPs-RhoB (red) accumulation in cultured microglial cells after incubation for 0.5 h, 1h, 2 h, 4 h or 24 h. Scale bar in D, 50 µm. F, Colocalization of NPs-RhoB (red) and ionized calcium-binding adaptor molecule 1 immunoreactivity (Iba1-ir, green) in microglial cells. G, No NPs-RhoB (red) was detected in astrocytes expressing the glial fibrillary acidic protein-ir (GFAP-ir, green). H, No NPs-RhoB (red) was detected in neurons expressing orexin (Green). n = 3. Scale bar: 20 μm in F, G, 30 μm in H.

### Effects of NPs-GW0742 on microglial activity and insulin sensitivity in diet-induced obesity

To assess the impact of NPs-GW0742 on food intake, body weight, and insulin sensitivity, we administered daily infusions of NPs-GW0742 into the MBH for 12 consecutive days, with blank NPs serving as controls (**Fig. 6A**). After the infusion period, no significant differences were observed in body weight or food intake between the NPs-GW0742 and blank NP groups (**Fig. 6B, C**). Basal blood glucose and insulin levels also remained unchanged (**Fig. 6D, E**). Interestingly, rats treated with NPs-GW0742 exhibited significantly improved insulin sensitivity during an insulin tolerance test (**Fig. 6F, G**). This improvement correlated with increased microglial activity, characterized by larger soma size and greater microglial cell coverage in the arcuate nucleus (ARC) of the MBH. Importantly, there was no difference in the total number of microglial cells between the groups (**Fig. 6H-K**). These results suggest that targeted activation of PPARδ signaling in microglia enhances their functional activity, thereby improving glucose metabolic regulation in diet-induced obesity.

**Figure 6.**
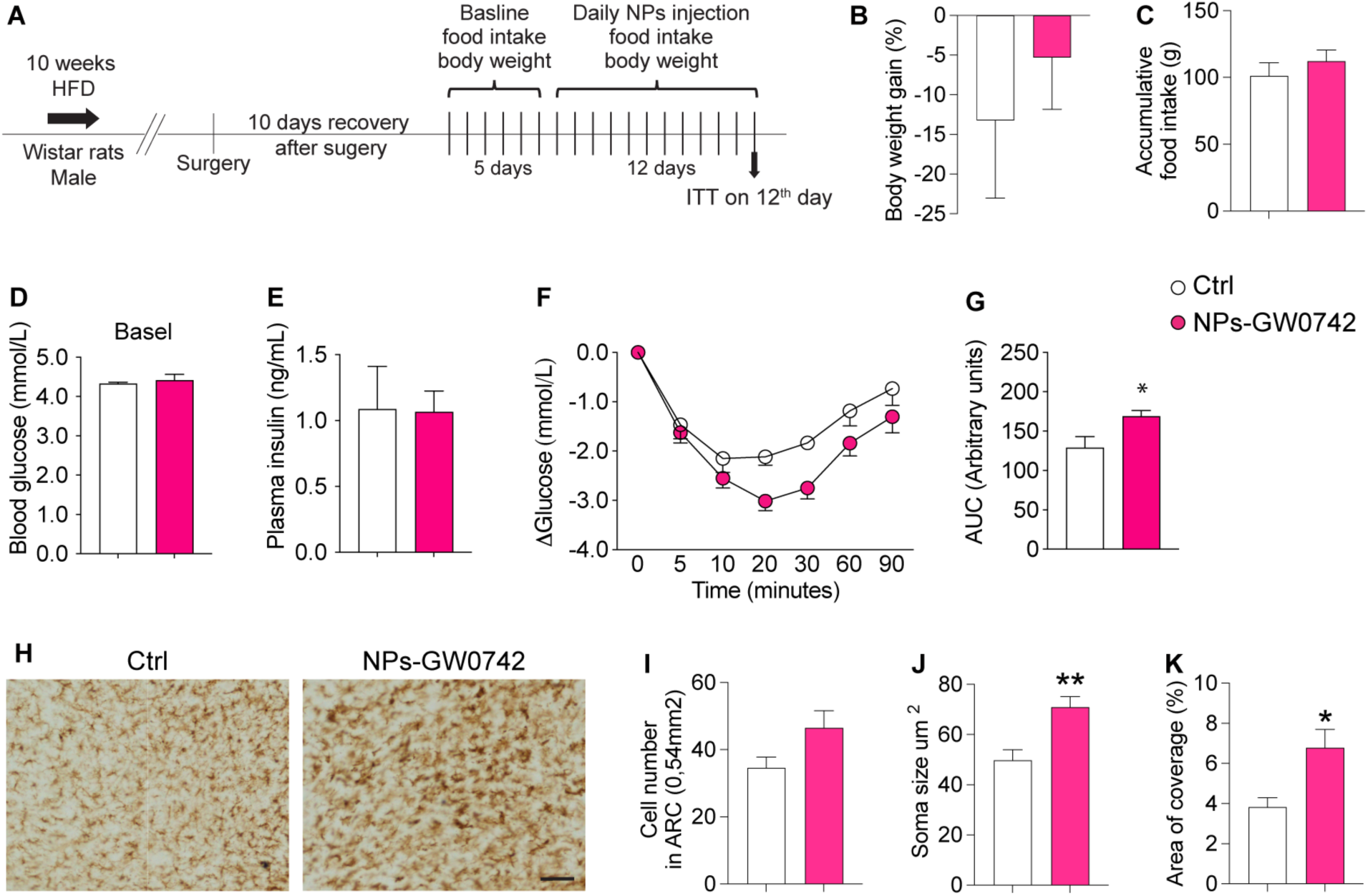
Effects of NPs-GW0742 Infusion into the mediobasal hypothalamus (MBH) on insulin sensitivity in rats with diet-induced obesity. A, Schematic overview of the in vivo experimental design. B, C, Body weight and food intake measured on day 12 following NPs-GW0742 infusion into the MBH. D, E, Blood glucose and plasma insulin levels on day 12 following NPs infusion. F, G, Blood glucose dynamics during the insulin tolerance test. H-K, Morphological changes in microglia, assessed by Iba1 immunoreactivity, following NPs-GW0742 treatment. Scale bar: 50 μm. Data are presented as means ± SEM, with statistical significance determined by an unpaired t-test.

## Discussion

This study highlights the critical role of microglial PPARδ activation as a novel approach to improving metabolic dysfunction associated with diet-induced obesity. Through the use of thermal proteome profiling, we identified GW0742 as a highly potent PPARδ ligand that modulates microglial function. The application of TPP not only provided insights into transcriptional regulation but also uncovered the impact of GW0742 on protein stability, with significant effects observed in mitochondrial and cytoskeletal proteins. These findings suggest that PPARδ activation in microglia induces an energy-efficient, anti-inflammatory state, which supports improved systemic insulin sensitivity.

The application of thermal proteomics in this study provided unique insights into the stability and functional state of microglial proteins following GW0742 treatment. Thermal proteome profiling enables the assessment of protein thermal stability, revealing potential changes in protein-protein interactions, ligand binding, and post-translational modifications [26]. This approach proved invaluable in elucidating how microglial metabolic pathways adapt to pharmacological modulation by PPARδ activation. Our proteomics analysis identified significant shifts in the thermal stability of proteins involved in mitochondrial function, energy metabolism, and inflammatory signaling. Notably, proteins associated with oxidative phosphorylation and mitochondrial biogenesis exhibited increased thermal stability post-GW0742 treatment. This suggests enhanced protein integrity and potential activation of mitochondrial pathways, aligning with the observed improvements in microglial metabolic capacity. The stabilization of these proteins may reflect an upregulation of their activity or an increased association with protective cofactors, which could facilitate more efficient energy production and better cellular resilience under metabolic stress.

The interaction between microglia and POMC neurons plays a crucial role in glucose metabolism and overall energy homeostasis. Previous studies from our group have demonstrated that microglia support POMC neurons through mechanisms such as maintaining metabolic equilibrium by regulating nutrient availability and releasing immune signals that influence neuronal activity [16, 17, 27, 33, 35, 42]. This regulation is essential for the proper maintenance of systemic glucose levels and body weight. Additionally, microglia contribute to the structural and functional integrity of POMC neurons by modulating synaptic pruning and plasticity, which is vital for effective signal transmission and neuropeptide release in response to metabolic cues [1, 23]. In the context of obesity, chronic microglial pro-inflammatory states disrupt POMC function and impair insulin signaling, leading to metabolic imbalances [42]. Our findings demonstrate that PPARδ activation through GW0742 induces an anti-inflammatory microglial state, reducing cytokine release and inflammation, potentially restoring POMC neuron function and improving glucose metabolism.

A key observation from our study was that chronic infusion of NPs-GW0742 into the MBH improved insulin sensitivity in diet-induced obese rats without affecting body weight or food intake. This finding, while unexpected, can be attributed to several factors. First, our experimental design only focused on delivering GW0742 specifically into microglial cells for 12 days into the MBH, we expect that longer treatment duration may be necessary to observe changes in body weight or feeding behavior. Second, the selective delivery of GW0742 to microglia may have limited its effects on broader hypothalamic networks that are involved in appetite and energy expenditure regulation. Future research should explore whether a more widespread activation of PPARδ across different cell types within the hypothalamus could yield more pronounced metabolic benefits. Additionally, the complex interplay of hormonal and neuronal signals that govern body weight likely means that while microglial activation improved insulin sensitivity, other regulatory pathways related to obesity may not have been sufficiently impacted by GW0742 in microglia.

The therapeutic implications of these findings are significant. Microglial-targeted therapies, specifically PPARδ agonists like GW0742, represent a promising avenue for addressing metabolic diseases, such as obesity and type 2 diabetes. The selective activation of microglial PPARδ not only shifts microglial immunometabolism towards an anti-inflammatory state but also preserves neuronal support functions that are disrupted in obesity. Future studies should investigate the long-term effects of microglial PPARδ activation on glucose homeostasis and metabolic health. Additionally, determining the optimal dosing and treatment duration for NPs-GW0742 will be critical for maximizing therapeutic efficacy. Expanding research to include other metabolic and neurodegenerative conditions characterized by microglial dysfunction and inflammation may reveal broader applications for PPARδ-targeted therapies. Moreover, exercise and diet are known to influence microglial activity and metabolism [16, 17, 27, 33, 35, 41, 42], potentially creating a synergistic effect with PPARδ activation. Therefore, a possible effective anti-obesity treatment combining GW0742 with lifestyle interventions, such as dietary changes and exercise [2, 7, 19], could further enhance metabolic outcomes. This combined approach could offer comprehensive treatment strategies that address both central and peripheral metabolic regulation.

### Experimental procedures

#### Cell culture

BV2 microglial cells were maintained in Dulbecco’s Modified Eagle Medium (DMEM; Gibco, 41965-039) supplemented with 10% fetal bovine serum (FBS) and 1% penicillin-streptomycin-neomycin (PSN). Cells were cultured in T-75 flasks at 37°C in a 5% CO₂ incubator until they reached confluence. Prior to experiments, cells were passage into 100 mm petri dishes and allowed to attach overnight. The medium was refreshed the next day to remove any cell debris. Treatments were performed when cells reached 80-90% confluency.

#### PPARδ agonists treatment

PPARδ agonists were prepared as 10 mM stock solutions in dimethyl sulfoxide (DMSO; Sigma-Aldrich, D8418-250ML) and stored at-20°C. On the day of the experiment, stocks were diluted to 2 mM in DMSO. The final working concentration of each agonist, including GNF-0242 (AKos GmbH, Z107275398), GNF-8501 (AKos GmbH, Z104661894), and GW-0742 (Sigma-Aldrich, G3295-5MG), was 1 μM in cell culture medium. DMSO at 0.05% (v/v) was used as a vehicle control in all treatments.

#### Lipopolysaccharide treatment

BV2 cells were exposed to 100 ng/mL LPS in serum-free Minimum Essential Medium (MEM) supplemented with PPARδ agonists for 24 hours. Serum-free MEM containing 0.05% DMSO served as the vehicle control. After treatment, cells were washed three times with phosphate-buffered saline (PBS) and collected for downstream analyses.

#### 2-NBDG glucose uptake assay

BV2 cells were seeded at a density of 1 × 10⁴ cells per well in 96-well black plates and treated with 1 μM PPARδ agonists or DMSO in complete culture medium for 6 hours at 37°C with 5% CO₂. Following treatment, cells were incubated with 200 µg/mL 2-NBDG (Abcam, ab235976), a fluorescently labeled deoxyglucose analog, and fluorescence was measured using a microplate reader with excitation and emission set at 480 nm and 530 nm, respectively.

#### Phagocytosis assay

Fluorescent latex microspheres (Sigma-Aldrich, L1030-1 ml) were pre-coated in PBS containing 10% FBS at 37°C for 1 hour. The microspheres were then centrifuged and re-suspended in DMEM with 0.05% FBS, reaching a final concentration of 0.01% (v/v) microspheres in the medium. BV2 cells were incubated with the microspheres for 1 hour, followed by three washes with cold PBS. Cells were fixed with 4% paraformaldehyde for visualization using confocal microscopy.

#### Measurement of microglial cellular respiration

BV2 cells (5000 cells/well) were plated in Seahorse XF96-well microplates and treated with 1 μM GW0742 or DMSO for 24 hours at 37°C with 5% CO₂. Prior to oxygen consumption rate (OCR) measurement, growth medium was replaced with Seahorse XF assay medium (basal DMEM containing 1 mM pyruvate, 2 mM glutamine, and 25 mM glucose), and cells were incubated in a non-CO₂ incubator for 1 hour. Baseline OCR was recorded over three cycles, followed by a mitochondrial stress test using sequential injections of oligomycin (1.5 μM), FCCP (0.5 μM), and a combination of rotenone (1.25 μM) and antimycin A (2.5 μM) as per the XF Cell Mito Stress Test protocol (Seahorse Bioscience). For extracellular acidification rate (ECAR) measurements, cells were incubated in glucose-and pyruvate-free DMEM with 1 μM GW0742 or DMSO. Glycolytic function was assessed using the XF Glycolysis Test kit (Agilent Technologies), with sequential injections of 10 mM d-glucose, 1 μM oligomycin, and 50 mM 2-deoxyglucose. Protein content for normalization was measured using the Bradford assay.

#### Real-Time PCR

Total RNA from BV2 cells was extracted using the RNeasy Micro Kit (QIAGEN, 74004) as per the manufacturer’s instructions. RNA concentrations were quantified using a NanoDrop Onec Spectrophotometer (Thermo Scientific, USA). cDNA synthesis was performed using HiScript II qRT Supermix (Vazyme Biotech, R222-01). Quantitative PCR was conducted on a QuantStudio 5 Real-Time PCR system (Applied Biosystems, Thermo Scientific) with AceQ qPCR SYBR Green Master Mix (Vazyme Biotech, Q131-02). Expression of target genes was normalized to housekeeping genes (Gapdh, Hprt, Rpl27). Primers were designed using NCBI BLAST and synthesized by GeneRay Biotech (Supplementary Table 1).

#### Sample preparation for Thermal Proteome Profiling

BV2 microglial cells were treated with 1 μM PPARδ agonists or DMSO in complete culture medium for 6 hours at 37°C with 5% CO2. After treatment, cells were detached by cell scraping and collected in 50 mL tubes. The cell suspension was centrifuged at 300g for 3 minutes to pellet the cells. The supernatant was discarded, and the cells were washed twice with 20 mL ice-cold PBS by centrifugation (300g, 3 minutes each). After washing, the supernatant was removed, and the cells were resuspended in 1 mL ice-cold PBS supplemented with protease inhibitors (Roche, 11873580001). Each treatment condition was divided into 8 Eppendorf tubes (500 µL per tube). The cells were centrifuged again at 300g for 3 minutes at 4°C, and the supernatant was carefully removed, leaving 20 µL of PBS in each tube. The samples were heated for 3’ in a Thermal Cycler at their designated temperature (37°C, 41°C, 46°C, 50°C, 54°C, 58°C, 62°C, 67°C). Immediately after heating the samples were incubated at room temperature (RT) for 3’, and then snap-frozen in liquid nitrogen.

The samples were prepared as presented previously with few modifications [12]. In short, thermally treated cell pellets were lysed three times with freeze-thaw cycles in 50 uL of 100 mM tri-ethyl-ammoniumhydrogencarbonate (TEAB), prior to clearing lysates of aggregated proteins and debris by centrifugation (21000g, 4°C, 1h). The protein concentration of the 37°C-treated cellular lysates were determined using the BCA Protein assay (23225, Thermo Scientific™) according to the manufacturer’s guidelines. Volumes equivalent to 20 ug (based on the BCA assay for the 37°C sample) were spiked with an internal standard (Bovine Serum Albumin), prior to reduction and alkylation with 10 mM tris-carboxy-ethyl-phosphine and 40 mM chloroacetamide for 15’ at 60°C. The samples were cooled to RT and cleaned up by Single-Pot Solid-Phase-enhanced Sample Preparation (SP3) as described previously [20]. The samples were subsequently digested with 1:20 (enzyme to substrate by weight) proteomics grade trypsin (V511, Promega) ON at 37°C. Peptide samples were subsequently cleaned by solid phase extraction using OASIS HLB extraction plates (WAT058951, Waters™) according to the manufacturer’s instructions and the resulting samples were dried in a vacuum centrifuge. Dried peptide samples were reconstituted in 1M TEAB for labelling using iTRAQ (8-plex kit, Thermo Fisher Scientific) according to the manufacturer’s guidelines. A small aliquot of each sample was checked for labelling efficiency before mixing and separating pooled TPP samples by strong cation exchange chromatography prior to mass spectrometric analysis.

#### Fractionation-Chromatography for Thermal Proteome Profiling

Pooled iTRAQ samples were separated by SCX chromatography to remove unreacted labels and fractionate peptides for deeper proteome coverage. For this, pooled iTRAQ samples were lyophilized and resuspended in solvent A (10 mM ammonium formate, 20% acetonitrile in ULCMS water, Biosolve) and injected onto a Polysulfoethyl aspartamide column (2.1 mm ID, 10 cm length, PolyLC Inc., Columbia, USA) with an Agilent 1100 HPLC (Agilent) fitted with a micro-fraction collector. Samples were separated by a linear gradient from 0% to 100 % solvent B (500 mM ammonium formate 20% acetonitrile in ULCMS water, Biosolve) and 1 min fractions were collected from 5-40 minutes. Fractions were subsequently pooled into 8 pools and lyophilized prior to mass spectrometric analysis.

#### Mass Spectrometry for Thermal Proteome Profiling

Dry peptide samples were reconstituted with 0.1% formic acid in water (232441, ULCMS grade, Biosolve) and 200 ng equivalent was injected onto a C18 column (75 µm, 250 mm, 1.6 µm particle size, Aurora, Ionopticks, Fitzroy, Australia) kept at 50°C by an on-source column oven (Sonation, Biberach, Germany). Using a NanoRSLC Ultimate 3000 UHPLC system (Thermo Scientific, Germeringen, Germany), peptide samples were separated by a multi-step gradient (Solvent A: 0.1% formic acid in water, Solvent B: 0.1% formic acid in acetonitrile). The peptides were loaded at 400 nl/min for 2 minutes in 3% solvent B and separated by a multi-step gradient, to 6% solvent B at 5 minutes, 21% solvent B at 21 minutes, 31% solvent B at 33 minutes, 42.5% solvent B at 36 minutes and 99% solvent B at 37 minutes held for 7 min before returning to initial conditions (Solvent A: 0.1% formic acid in water, Solvent B: 0.1% formic acid in acetonitrile) until 60 minutes (total run time). The eluting peptides were electro-sprayed by a captive-spray source, into a timsTOF-pro (trapped ion mobility spectroscopy, quadrupole time of flight mass spectrometer, Bruker, Bremen, Germany), using the following settings: precursor scan ranged from 100 to 1700 m/z and a tims range of 0.6–1.6 V.s/cm2 in PASEF mode. A total of 10 PASEF MS/MS scans were collected with a total cycle time of 1.16 s. All further settings were as described previously by Ogata et al. (2021) for optimized detection of isobaric labels [29].

#### Thermal Proteome Profiling data analysis

Resulting mass spectra were analyzed by Maxquant (v.1.6.10.43), searching against the proteome database of Mus musculus (Uniprot, 10/2019, https://www.uniprot.org). Search parameters were set for isobaric labelling (iTRAQ 8-plex, using the isotopic correction factors provided with the kit), with the enzyme set as trypsin, allowing for 2 missed cleavages, and carbamidomethylation at cysteine as a fixed modification, and oxidation at methionine and n-terminal acetylation as variable modifications. Protein false discovery rate was kept at 1% using a target-decoy database. Identified protein groups were further processed using R (v.4.1.2), with the Bioconductor package TPP [8], to ascertain melting curves and differences in melting temperatures between treatment and vehicles, with parameters adjusted to accommodate the iTRAQ 8-plex labelling kit [8].

#### Synthesis of blank NPs and NPs-GW0742

In this study, we synthesized biocompatible and biodegradable methoxy poly(ethyleneglycol)-poly(lactide-co-glycolide) (mPEG-PLGA) nanoparticles (NPs) for use as drug carriers (Poly (lactic-co-glycolic acid) (PLGA, molar ratio of D, L-lactic to glycolic acid, 75:25), monomethoxy poly (ethylene glycol) (mPEG) purchased from Jinan Daigang Biotechnology Co. Ltd.). These NPs have garnered significant attention due to their advantageous physicochemical properties and safety profiles [18]. The incorporation of a poly(ethyleneglycol) (PEG) coating enhances the solubility and stability of the NPs within the brain microenvironment. Specifically, NPs-GW0742 were fabricated through an emulsion/solvent evaporation method. A mixture of 5 mg of copolymer and 0.1 mg of GW0742 was dissolved in 1 mL of dichloromethane. The solution was stirred for 10 minutes at room temperature and subsequently emulsified via sonication in 10 mL of aqueous solution containing 0.6% PVA. Following the vacuum evaporation of the solvent, the NPs were collected through centrifugation at 13,000 rpm for 10 minutes at room temperature and were then washed twice with distilled water.

*Characterization of Blank NPs and NPs-GW0742:* The size (diameter in nm), polydispersity index, and surface charge (zeta potential, mV) of the nanoparticles were assessed using a Zeta Sizer Nano ZS (Malvern Instruments Ltd., Malvern, UK), which is equipped with a HeNe laser operating at a wavelength of 633 nm and a fixed scattering angle of 90°. Measurements were conducted at 25 °C using samples diluted in distilled water. The morphology of the nanoparticles was analyzed via transmission electron microscopy (TEM, JEM-200CX, Jeol Ltd., Japan) following negative staining with a 2% (w/w) sodium phosphotungstate solution.

*Stability, encapsulation efficiency and drug release of NPs-GW0742 in vitro:* The blank NPs (50 mg) were suspended in phosphate-buffered saline (PBS) at pH 7.4 and pH 4.4, and the suspension was stirred at 110 rpm at 37 °C. The size of the nanoparticles was measured at 0, 1, 2, and 3 days using dynamic light scattering (DLS) to evaluate the stability of the nanoparticles at different pH levels.

To evaluate the encapsulation of GW0742 in the polymeric NPs, the NPs-GW0742 were treated with dichloromethane (DCM) to break the mPEG-PLGA shell and release the encapsulated cargo. Following DCM removal by rotary evaporation, the residual GW0742 was dissolved in acetonitrile. The concentration of GW0742 was determined using high-performance liquid chromatography (HPLC) (SPD-20A/20AV Series equipped with a SIL-20A/20AC detector and a Thermo Scientific™ C18 reversed-phase column, Part No. 25005-154630), with detection at a wavelength of 226 nm. Standard curves were constructed using dilutions of GW0742 at various concentrations. The encapsulation efficiency (EE) was calculated using the following equation: Encapsulation Efficiency (EE) = (B-A)/B *100%, where A represents the amount of drug released into the supernatant and B denotes the initial amount of drug added to the system.

#### Vessel cannulation and brain infusion probe implantation

All animal procedures were conducted in accordance with the guidelines on animal experimentation of the Netherlands Institute for Neuroscience (NIN, Amsterdam) and were approved by the Animal Ethics Committee of the Royal Dutch Academy of Arts and Sciences (KNAW; Amsterdam). Male Wistar rats (Charles River, Germany), aged 8-10 weeks (body weight: 150–200 g), were housed under a 12-hour light/12-hour dark cycle (lights on at 7:00 am) at a controlled temperature of 22 ± 2°C, with unrestricted access to food and water. All rats were fed a HFD (Research Diet, D12331, 58 kcal% Fat and Sucrose) for 10 weeks before undergoing surgery. For vessel cannulation and brain infusion probe implantation, rats were anesthetized with an intramuscular injection of ketamine (80 mg/kg; Eurovet Animal Health, Bladel, Netherlands) combined with xylazine (8 mg/kg; Rompun, Bayer Health Care, Mijdrecht, Netherlands). Anesthesia was monitored throughout the procedure to ensure depth and stability.

A standard Kopf stereotaxic apparatus was used to ensure precise placement of the brain infusion probes. For bilateral injections of RhoB-labeled nanoparticles (NPs-RhoB) into the mediobasal hypothalamus, a Hamilton microliter syringe (7000 series; Hamilton) was utilized to inject 1 μl of solution per side. Coordinates were adapted from the Paxinos and Watson atlas: anteroposterior −2.8 mm, lateral ±2.0 mm, angle 8°, and ventral −10 mm. Rats were euthanized 24 hours post-infusion via perfusion fixation with 0.9% saline followed by 4% paraformaldehyde. Brains were collected and post-fixed in 4% paraformaldehyde for 16 hours prior to immunohistochemical and immunofluorescent analysis.

#### Chronic Infusion and Blood Sampling

For long-term delivery of NPs and concurrent blood sampling for the insulin tolerance test, silicone catheters were inserted into the right jugular vein and left carotid artery. These catheters facilitated intravenous infusions and blood collection, respectively. Bilateral micro-infusion guide cannulas (P1 Technologies) were also implanted into the mediobasal hypothalamus, with coordinates: anteroposterior:-3.3 mm; lateral: 2.0 mm; ventral:-9.7 mm; angle: 8°), and secured using dental cement. Post-surgical recovery was monitored for up to 10 days. Rats were included in the study only if they regained their pre-surgical body weight within this period; those that did not meet this criterion were excluded. For chronic administration, an infusion probe was inserted into the implanted guide cannula, and 0.5 μl of blank NPs or NPs-GW0742 (1μM of GW0742) was delivered at a controlled rate using a Hamilton microliter syringe. The infusion rate was set at 0.1 μl per click, with one click per minute, continuing for a total of 12 days. Food intake, body weight, and behavior were monitored daily during the infusion period. On the final day, rats were sacrificed by perfusion fixation, and brains were collected for subsequent immunohistochemical or immunofluorescent analyses.

#### Insulin tolerance test

To evaluate the impact of NPs-GW0742 on whole body glucose metabolism, in a separate batch of rats, following 12 days of nanoparticle (NP) infusion, an insulin tolerance test (ITT) was performed. To minimize stress, rats were acclimated to handling during infusions. Insulin administration and blood sampling (200 μL per sample) were conducted in the rats’ home cages to allow free movement. After each blood sample, an equivalent volume of saline was administered to maintain fluid balance. A baseline blood sample (t = 0) was collected before the intravenous administration of an insulin bolus (0.1 IU/kg body weight; Actrapid, Novo Nordisk, Bagsværd, Denmark). Subsequent blood samples were collected at t = 5, 10, 20, 30, 60, and 90 minutes post-injection to monitor glucose levels. At the conclusion of the experiment (t = 90 minutes), rats were anesthetized using pentobarbital and euthanized via decapitation.

#### Immunohistochemical and immunofluorescent staining

For morphological characterization, rats that received NPs infusions were anesthetized with pentobarbital and underwent intra-atrial perfusion with saline, followed by 4% paraformaldehyde in 0.1 mol/L phosphate buffer (pH 7.4) at 4°C. Brains were post-fixed in the same paraformaldehyde solution for 16 hours and subsequently equilibrated in 30% sucrose. Coronal sections (35 µm thick) were prepared using a cryostat. From each brain, 4–6 sections encompassing the injection sites (2–3 sections per side) were selected, as the diffusion area of the nanoparticles is approximately 300 μm from the injection site [18]. Sections exhibiting mechanical damage from the infusion needle or brain probes were excluded, and only the highest quality sections were used for staining.

To detect NP-RhoB in the brain, immunofluorescent staining was conducted on selected sections. Two sections per brain were stained for Iba1, two consecutive sections for GFAP, and another two consecutive sections for orexin neurons in the lateral hypothalamus. The sections were incubated overnight at 4°C with the following primary antibodies: Iba1 (Synaptic Systems, No. 234003; 1:400), GFAP (DAKO, Z0334; 1:400), and orexin (Abcam, Ab6214; 1:1000). Secondary antibodies (Vector Lab, BA1100; 1:400) and streptavidin Alexa Fluorophore 488 (Invitrogen, s32354; 1:300) were applied for 1 hour. After staining, sections were dried and mounted with Antifade Mounting Media (VECTASHIELD; Vector Labs). Fluorescent signals were visualized using a Leica TCS SP8 confocal microscope.

For evaluate the impact of NPs-GW0742 on microglia, single immunohistochemical analysis of Iba1 was performed. Two to three sections per brain for each rat were incubated overnight at 4°C with Iba1 primary antibody (Synaptic Systems, No. 234003; 1:2000). Sections were rinsed and incubated with a biotinylated secondary antibody (Vector Lab, BA1100; 1:400) and avidin-biotin complex (Vector Laboratories, Inc., 1:800). The reaction was visualized with 0.5% diaminobenzidine and 0.01% hydrogen peroxide. Sections were mounted, dehydrated in ethanol, cleared in xylene, and covered with Entelan-mounted coverslips. Images were captured using a Zeiss Axioplan 2 Evolution MP camera. Analysis of cell count, soma size, and coverage area within the arcuate nucleus (ARC) in the mediobasal hypothalamus was performed using FiJi software, with data averaged per rat for statistical evaluation.

## Statistical Analysis

All the results were presented as means ± SEM. Two tailed Student t’s-test, one-way or two-way ANOVA followed by post hoc analysis was used to test for differences between individual experimental groups. Differences between groups were considered statistically significant when two-sided p < 0.05. Statistics were calculated using IBM SPSS version 22 or GraphPad Prism 7 software.

## Author contributions

H.J. performed cell treatments with compounds for in vitro profiling and Thermal proteome profiling, G.K. performed Thermal proteome profiling and data analysis, S.G. performed nanoparticle synthesis, characterization and functional analysis by *in vitro* experiments, S.G., F.C.M. performed the *in vivo* studies. H.J., F.C.M., S.G., G.K., A.K. and C.X.Y. wrote the manuscript. S.G., G.K. and C.X.Y conceived the idea and the experimental design. All co-authors discussed the results and commented on the manuscript.

## Funding and additional information

This study was supported by ZonMw Enabling Technologies Hotels 5^th^ programme 2020 (435005036), The Netherlands.

---

**Supplementary table 1.**
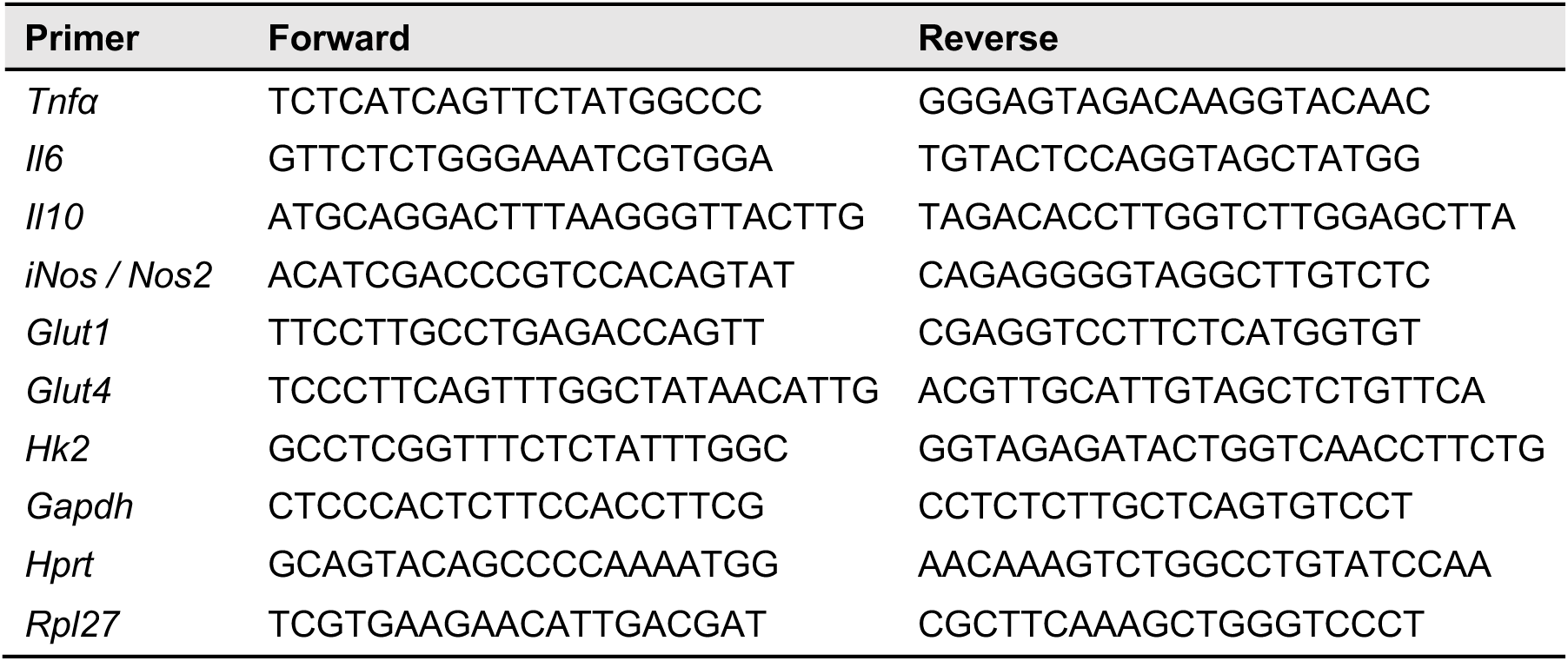
Primer sequences.

